# Immunomodulatory Leptin Receptor^+^ Sympathetic Perineurial Cells Protect Against Obesity by Facilitating Neuroendocrine-Mediated Brown Adipose Tissue Thermogenesis

**DOI:** 10.1101/2023.05.09.539963

**Authors:** E R Haberman, G Sarker, B A Arús, S Yilmaz-Özcan, N Martínez-Sánchez, E Freibergerová, I Fernández-González, C Zentai, C J O O’Brien, D E Grainger, S Chakarov, A Raimondi, M Iannacone, M López, F Ginhoux, A I Domingos

## Abstract

Adipose tissues (ATs) are innervated by sympathetic nerves, which drive reduction of fat mass via lipolysis and thermogenesis. Here, we report a population of immunomodulatory leptin receptor (LepR)-expressing barrier cells which ensheath sympathetic axon bundles in adipose tissues. These LepR-expressing Sympathetic Perineurial Cells (SPCs) produce IL33, a factor for maintenance and recruitment of regulatory T cell (Treg) and eosinophils in AT. Brown adipose tissues (BAT) of mice lacking IL33 in SPCs (SPC^IL33cKO^) have fewer Treg and eosinophils, resulting in increased BAT inflammation. SPC^IL33cKO^ mice are more susceptible to diet-induced obesity, independently of food intake. Furthermore, SPC^IL33cKO^ mice have impaired adaptive thermogenesis, and are unresponsive to leptin-induced rescue of metabolic adaptation. We, therefore, identify LepR-expressing SPCs as a source of IL33 which orchestrate an anti-inflammatory environment in BAT, preserving sympathetic-mediated thermogenesis and body weight homeostasis. LepR^+^ IL33^+^ SPCs provide a cellular link between leptin and immune regulation of body weight, unifying neuroendocrinology and immunometabolism as previously disconnected fields of obesity research.

**Graphical Abstract:** 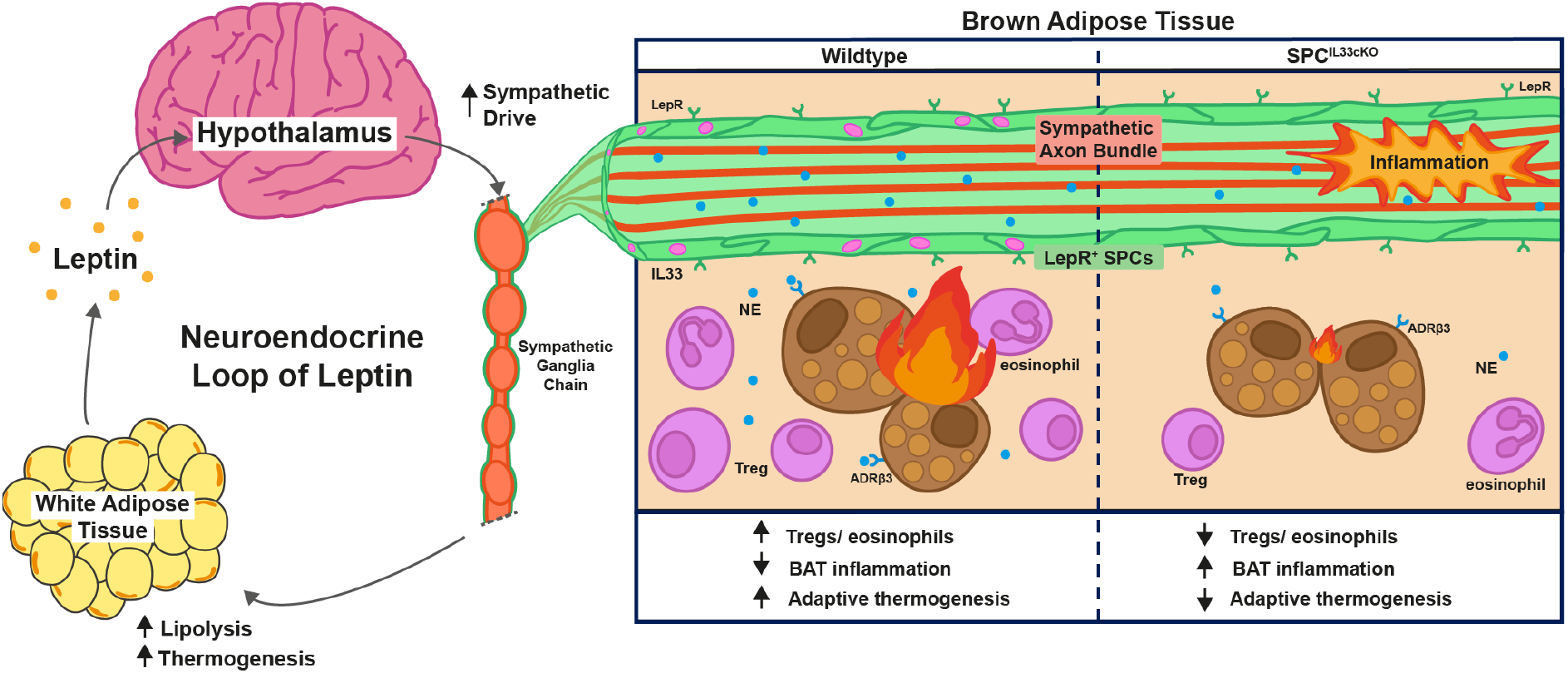

**Highlights:**

- Sympathetic Perineurial Cells (SPCs) co-express LepR^+^ and IL33
- SPC-derived IL33 prevents BAT inflammation via Treg and eosinophil recruitment
- Obesity is worsened in high fat diet-fed SPC^IL33cKO^ mice, despite normal food intake
- Adaptive thermogenesis is impaired in SPC^IL33cKO^ mice
- Rescue of metabolic adaptation to fasting by leptin is impaired in SPC^IL33cKO^ mice
- SPCs link leptin to immunometabolic regulation of body weight homeostasis

## Introduction

Adipose tissue (AT)-resident immune cells have been extensively implicated in obesity, which is linked with chronic, low-grade inflammation of the adipose tissue^1^. This is reflected in an increase of pro-inflammatory adipose tissue macrophages and a concomitant reduction in anti-inflammatory immune cells, including regulatory T cells (Tregs), and eosinophils^2–8^. In tandem with immunometabolic regulation, body weight is chiefly controlled by the hormone leptin. Leptin is released from ATs proportional to fat mass where it acts on hypothalamic leptin receptor-positive (LepR^+^) neurons to increase downstream sympathetic drive onto adipose tissues, stimulating thermogenesis in brown AT (BAT)^1,9–11^. This neuroendocrine loop of leptin action is disrupted in obesity, where sympathetic drive onto ATs is suppressed and BAT thermogenesis is blunted, further worsening weight gain^12^. Studies on the neuroendocrine and immune regulation of adipose tissues have been separated, with no biological players linking the two fields^13^. Here we identify one such cellular intermediate, providing a missing link bridging the neuroendocrine loop of leptin action with immune regulation of energy homeostasis.

## Results

### A LepR^+^ Sympathetic Perineurial Cell (SPC) barrier ensheathes sympathetic ganglia and axon bundles

Sympathetic neurons innervate adipose tissues via the paravertebral sympathetic ganglia (Figure 1A). Considering the role of leptin on AT lipolysis and thermogenesis^10^, we investigated the presence of leptin receptor^+^ (LepR^+^) cells within AT sympathetic axon bundles. Immunofluorescent (IF) imaging of LepR^eYFP^ sympathetic tissues revealed a LepR^+^ cell barrier ensheathing both subcutaneous white AT (scWAT) and BAT axon bundles (Figure 1B-D), as well as sympathetic ganglia (Supplementary Figure 1A). We detected similar frequencies of LepR^+^ cells in all sympathetic tissues (Supplementary Figure 1B-C). To visualise the ultrastructure of this LepR^+^ cell barrier, we performed immuno-electron microscopy (immuno-EM) on cross sections of LepR^eYFP^ scWAT axon bundles, using gold-conjugated anti-eYFP antibodies to mark LepR^+^ cells. We saw that the LepR^+^ barrier surrounding sympathetic AT nerves is formed of multiple, thin LepR^+^ cells (Figure 1E) – similar to the perineurium^14–16^.

**Figure 1:**
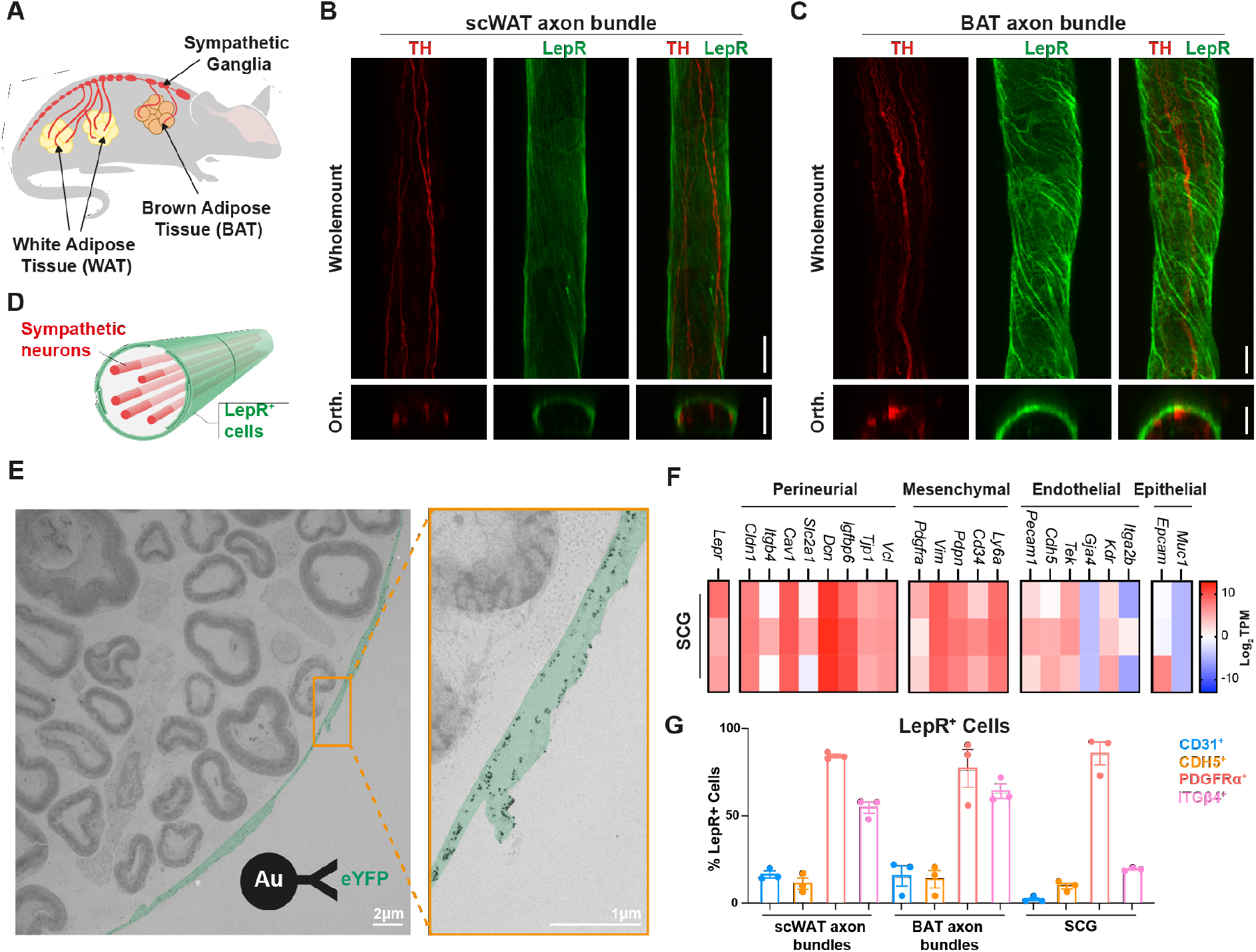
A LepR^+^ Sympathetic Perineurial Cell (SPC) barrier ensheathes sympathetic ganglia and axon bundles. **A)** Schematic representation: adipose tissue is innervated by sympathetic axon bundles originating from the paravertebral chain of sympathetic ganglia. ^10,40,41^ **B-C)** Sympathetic axon bundles dissected from the subcutaneous white adipose tissue (scWAT), or brown adipose tissue (BAT) are surrounded by a LepR^+^ cell barrier. LepR^eYFP^ mice, 20x maximum intensity projection images (Wholemount) or orthogonal optical reconstruction (Orth). TH - tyrosine hydroxylase (red), LepR^eYFP^ (green). Scale bars = 25μm. **D)** Schematic representation: sympathetic neurons are surrounded by a LepR^+^ barrier in adipose tissue nerves. **E)** Transmission electron micrograph of a scWAT sympathetic axon bundle cross section. Immuno-EM staining with gold-conjugated anti-eYFP antibodies reveal a multi-layer LepR^+^ perineurial cell barrier. scWAT fibre dissected from LepR^eYFP^ mice. LepR cells false-coloured in green. Scale bars = 2μm. **F)** Bulk RNA-sequencing of LepR^+^ cells sorted from the Superior Cervical Ganglia (SCG) of LepR^eYFP^ mice (n=3). LepR^+^ cells highly express perineurial and mesenchymal cell markers. Data presented as Log_2_ transcripts per million (TPM). **G)** Flow cytometric analysis of LepR^+^ cells isolated from sympathetic tissues (scWAT axon bundles, BAT axon bundles and SCG). LepR^+^ Sympathetic Perineurial Cells (SPCs) in sympathetic tissues are PDGFRα^+^ITGβ4^+^. LepR^eYFP^ mice (n=3). Full gating strategy in Supplementary Figure 1B. Data presented as Mean+/-SEM.

To resolve their identity, we sorted LepR^+^ cells from the Superior Cervical Ganglia (SCG) of LepR^eYFP^ mice and performed bulk RNA sequencing. Gene expression analysis revealed that these LepR^+^ cells express a range of perineurial (*Cldn1, Cav1, Dcn, Igfbp6, Tjp1, Vcl*) and mesenchymal markers (*Pdgfra, Vim, Pdpn, Cd34, Ly6a*), with low expression of endothelial markers (*Pecam1, Cdh5, Tek, Kdr*) and an absence of epithelial marker (*Epcam, Muc1*) expression (Figure 1F). Using flow cytometry, we verified at the protein level that the vast majority of LepR^+^ cells in sympathetic tissues stained positive for fibroblastic and perineurial markers, PDGFRα and ITGβ4, respectively (Figure 1G). Few LepR^+^ cells were positive for endothelial markers CD31 and CDH5 (Figure 1G). Based on this characterisation, we name these specialised LepR^+^ cells Sympathetic Perineurial Cells (SPCs).

### LepR^+^ SPCs produce anti-inflammatory IL33

Having characterised LepR^+^ SPCs, we hypothesised SPCs have an immunometabolic role, preventing the chronic, low-grade AT inflammation closely associated with obesity. Analysis of our RNA-sequencing dataset revealed high expression of anti-inflammatory cytokines Interleukin 33 (*Il33*), Transforming growth factor β2 (*Tgfb2*), and Interleukin 4 (*Il4*) in LepR^+^ SPCs (Figure 2A). Recent studies have implicated IL33 and IL33-responsive immune cells in the control of AT homeostasis^17–19^. Using flow cytometry, we found that the majority of LepR^+^ SPCs in all sympathetic tissues are IL33^+^ (Figure 2B-C) and verified the presence of IL33 protein in scWAT and BAT axon bundles using IF imaging (Figure 2D-E). Orthogonal, optical reconstructions of wholemount-imaged scWAT and BAT axon bundles show exclusive IL33 staining in the outer LepR^eYFP+^ perineurial barrier cells (Figure 2D-E).

**Figure 2:**
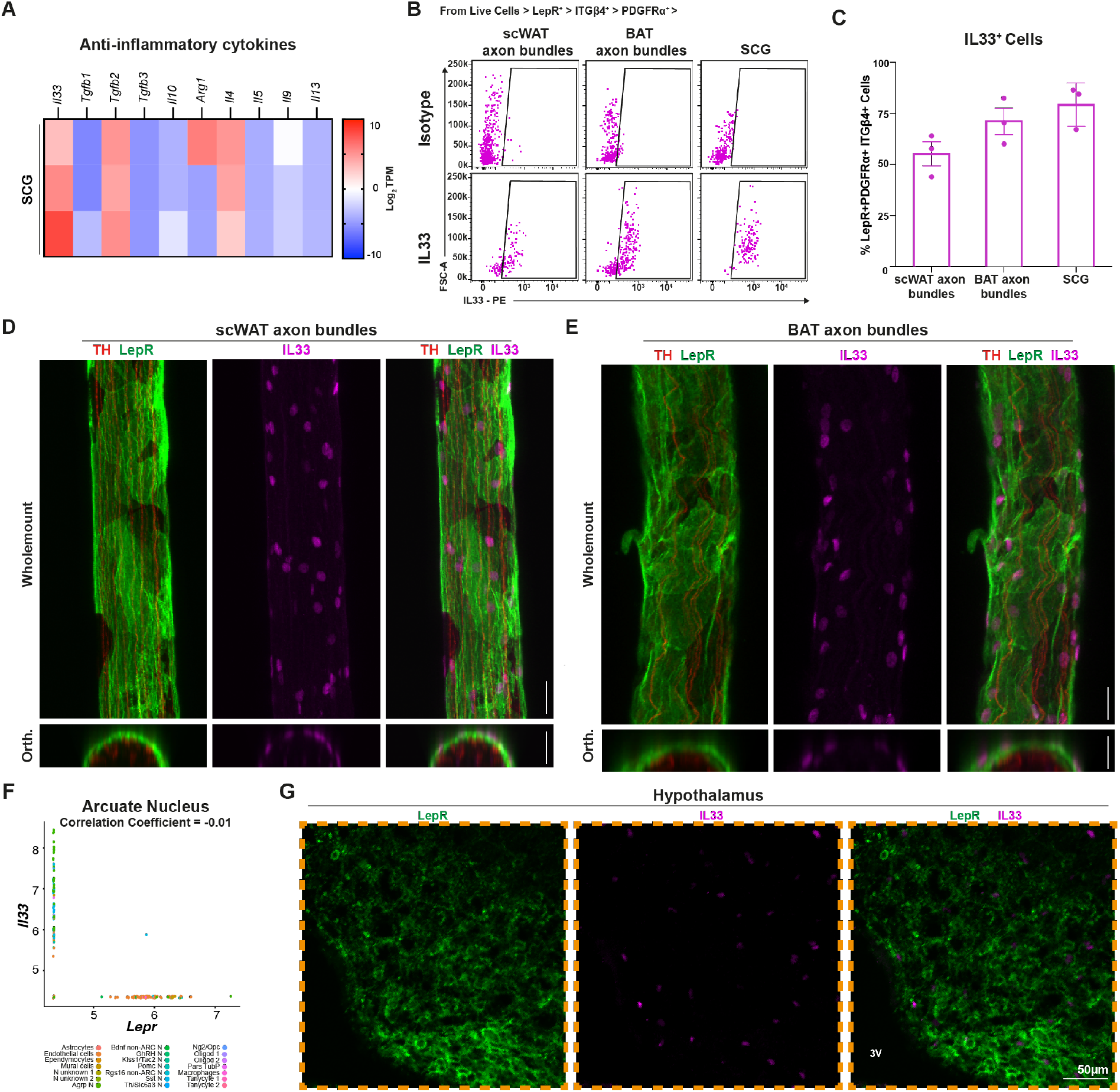
LepR^+^ Sympathetic-Associated Perineurial Cells (SPCs) produce anti-inflammatory IL33. (A) Bulk RNA-sequencing of LepR^+^ cells sorted from the Superior Cervical Ganglia (SCG) of LepR^eYFP^ mice shows that LepR^+^ cells express anti-inflammatory cytokines, including *Il33*. n=3, data presented as Log_2_ transcripts per million (TPM). **B-C)** Flow cytometric analysis reveals that the majority of LepR^+^ SPCs (PDGFRα^+^ITGβ4^+^) in sympathetic tissues are IL33^+^. LepR^eYFP^ mice (n=3). Full gating strategy in Supplementary Figure 1B. **D-E)** LepR^+^ Sympathetic Perineurial Cells (SPCs) in subcutaneous (scWAT, **D**) and brown (BAT, **E**) adipose tissue axon bundles are IL33^+^. LepR^eYFP^ mouse, 20× maximum intensity projection images (Wholemount) or orthogonal reconstruction (Orth). TH - tyrosine hydroxylase (red), LepR^eYFP^ (green), IL33 (magenta). Scale bars = 25μm. Data presented as Mean+/-SEM. **F)** Cells within the arcuate nucleus of the hypothalamus (ARC) do not co-express *Lepr* and *Il33* (correlation coefficient = -0.01). Analysis of mouse ARC single-cell RNA sequencing dataset ^20^. N = neuron, Oligod = oligodendrocytes, Pars Tub = Pars Tuberalis. **G)** Immunofluorescent staining of Coronal LepR^eYFP^ hypothalamus section, maximum intensity projection image (10x). LepR^eYFP^ (green), IL33 (magenta). Anatomical location of image shown with orange box in Supplementary Figure 2A. 3V = third ventricle. Scale bar = 50μm.

### *Lepr* and *Il33* co-expression is not observed in the hypothalamus, or organs of the *Tabula Muris* database

To exclude the possibility of off-target effects for a LepR^cre^-mediated conditional IL33 knockout mouse model, we first assessed whether any other cell in wildtype mice is double positive for *Lepr* and *Il33*. We analysed public single-cell RNA sequencing data from the arcuate nucleus of the hypothalamus (ARC)^20^ and found that cells in the ARC do not co-express *Lepr* and *Il33* (Figure 2F), which we confirmed using immunofluorescent imaging (Figure 2G, Supplementary Figure 2A). We next assessed *Lepr* and *Il33* co-expression in peripheral organs using the *Tabula Muris* single-cell RNA sequencing database^21^, which revealed that the kidney, muscle and trachea were the most likely to co-express *Lepr*/*Il33*, with correlation coefficients between 0.1–0.4 (Supplementary Figure 2B). However, immunofluorescent imaging excluded the possibility of LepR^+^IL33^+^ cells in these organs (Supplementary Figure 2C). Using immunofluorescent imaging we also excluded *Lepr* and *Il33* co-expression in the aorta, heart, and liver, with co-expression correlation coefficients of 0.08-0.1 (Supplementary Figure 2C). The diaphragm, pancreas, tongue, bone marrow, skin, colon, thymus, and bladder all had very low *Lepr*/*Il33* correlation coefficients, below 0.05, and hence LepR^+^IL33^+^ cells in these organs can be safely excluded (Supplementary Figure 2B). We, therefore, confirm that LepR^+^IL33^+^ expressing cells are not present in the hypothalamus, nor in the peripheral tissues listed in the *Tabula Muris* database.

### Loss of IL33 in LepR^+^ SPCs drives BAT inflammation

To functionally assess the role of IL33 on SPC cell function, we crossed LepR^cre^ mice with IL33^fl/fl^ mice to perform a conditional knockout of IL33 only in SPCs (herein named SPC^IL33cKO^ mice). Having validated that, other than the SPCs, no cells in the hypothalamus or organs of the *Tabula Muris* co-express *Lepr* and *Il33*, we next verified using IF that IL33 is lost in SPCs in SPC^IL33cKO^ AT axon bundles (Figure 3A) and found that lean SPC^IL33cKO^ mice fed a normal diet have comparable body weights, adipose tissue weights and food intake to IL33^fl/fl^ controls (Figure 3C-G, Supplementary Figure 3A-B). Considering the role of IL33 in the type 2 immune response, we next tested if known IL33-responsive (ST2^+^) adipose tissue immune cells – including regulatory T cells (Tregs) and eosinophils – were reliant on SPC-derived IL33 ^22–29^. Whilst the spleen and white adipose tissue (WAT) depots appeared unaffected, CD45^+^ cell count was significantly reduced in the BAT of SPC^IL33cKO^ mice (Figure 3H). Upon closer inspection, this was due to a decrease in the frequency of both BAT Tregs and BAT eosinophils (Figure 3I-J), with no significant differences in splenic or scWAT Treg and eosinophil frequencies (Supplementary Figure 3C-F). Whilst visceral white adipose tissue (vWAT) Treg frequency remained unchanged, a significant decrease in vWAT eosinophils was observed in SPC^IL33cKO^ mice (Supplementary Figure 3D-E). Specifically, IL33-responsive immune cells seem to be most significantly affected in the brown adipose tissue of SPC^IL33cKO^ mice. In addition to changes in BAT-resident immune cells, we observed that the expression of *Il1b* was increased in SPC^IL33cKO^, indicating some BAT inflammation in lean SPC^IL33cKO^ mice (Figure 3K). In summary, we show SPC^IL33cKO^ manifests in significant changes to BAT immune cell recruitment and maintenance.

**Figure 3:**
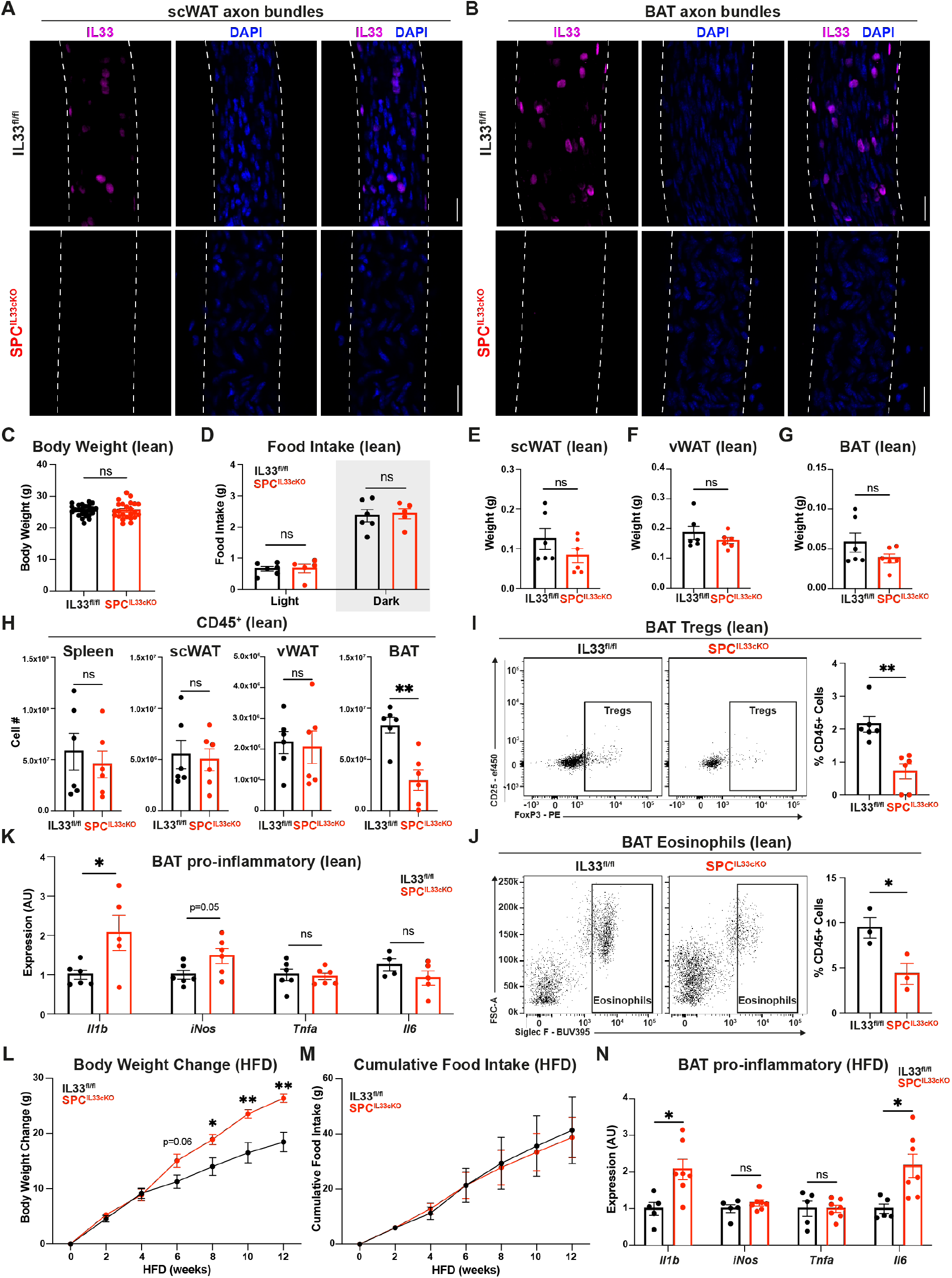
Loss of IL33 in LepR+ SPCs induces brown adipose tissue inflammation and predisposes mice to obesity, independently of food intake. **A-B)** IL33 is lost in the LepR^+^ Sympathetic Perineurial Cells (SPCs) of SPC^IL33cKO^ subcutaneous (scWAT, **A**) and brown (BAT, **B**) adipose tissue axon bundles. IL33^fl/fl^ and SPC^IL33cKO^ mice, 20× maximum intensity projection images. IL33 (magenta), DAPI (blue). Scale bars = 25μm. White dotted lines mark nerve edges. **C)** Absolute body weight of lean (normal diet-fed) IL33^fl/fl^ and SPC^IL33cKO^ mice. n=24, 3 independent experiments. **D)** Average food intake per mouse of normal diet-fed IL33^fl/fl^ and SPC^IL33cKO^ mice during light/dark periods, measured over 5 days. n=5-6, 1 independent experiment. **E-G)** Comparable adipose tissue weights between lean (normal diet-fed) IL33^fl/fl^ and SPC^IL33cKO^ mice. Subcutaneous white adipose tissue (scWAT, **E**), visceral white adipose tissue (vWAT, **F**) and brown adipose tissue (BAT, **G**). n=6, 2 independent experiments. **H)** Absolute number of CD45^+^ cells in the spleen, subcutaneous (scWAT), visceral (vWAT) and brown adipose tissue (BAT) of IL33^fl/fl^ and SPC^IL33cKO^ mice. n=6, 2 independent experiments. **I-J)** Flow cytometric analysis of brown adipose tissue (BAT) from lean (normal diet-fed) IL33^fl/fl^ and SPC^IL33cKO^ mice shows a reduction in the frequency of BAT Tregs (**I**) and eosinophils (**J**). **I**) n=6, 2 independent experiments. **J**) n=3, 1 independent experiment. **K)** qPCR of BAT shows increased inflammation in lean (normal diet-fed) SPC^IL33cKO^ mice. For each gene, data is presented relative to lean IL33^fl/fl^ expression level (=1). *Il1b* = Interleukin-1 beta, *iNos* = Inducible nitric oxide synthase, *Tnfa* = Tumor necrosis factor alpha, *Il6* = Interleukin-6. n=4-6, 2 independent experiments. **L-M)** SPC^IL33cKO^ mice gain more weight than IL33^fl/fl^ controls when fed high-fat diet (HFD, **L**), despite no difference in food intake (**M**). Body weight change (**L**) of IL33^fl/fl^ and SPC^IL33cKO^ mice fed HFD from 8wo. n=6, 2 independent experiments. **N)** HFD-fed SPC^IL33cKO^ mice have increased BAT inflammation compared with HFD-fed IL33^fl/fl^ controls. qPCR shows higher expression of pro-inflammatory genes *Il1b* and *Il6* in SPC^IL33cKO^ BAT. For each gene, data is presented relative to IL33^fl/fl^ HFD expression level (=1). *Il1b* = Interleukin-1 beta, *iNos* = Inducible nitric oxide synthase, *Tnfa* = Tumor necrosis factor beta, *Il6* = Interleukin-6. n=5-7, 1 independent experiment. Data presented as Mean+/- SEM. Student’s t-test, ns = non-significant, *p<0.05, **p<0.01.

### Production of IL33 by LepR^+^ SPCs is protective against diet-induced obesity

Although SPC^IL33cKO^ mice did not develop obesity when fed a normal diet, SPC^IL33cKO^ mice metabolically challenged with a high-fat diet (HFD) from 8 weeks old for 12 weeks gained more weight independently of food intake, relative to age-matched, HFD-fed IL33^fl/fl^ controls (Figure 3L-M, Supplementary Figure 3G). Furthermore, HFD-fed SPC^IL33cKO^ mice had increased scWAT weights than control mice, although no significant difference was observed in vWAT or BAT weights (Supplementary Figure 3H-J). Having identified the BAT as a potential site of metabolic dysregulation in SPC^IL33cKO^ mice (Figure 3I-J), we also found using qPCR that HFD-fed SPC^IL33cKO^ BAT was more inflamed than HFD-fed control BAT, with increased expression of *Il1b* and *Il6* (Figure 3N). Collectively, this further supports a role for SPC-derived IL33 in BAT function.

### IL33 derived from LepR^+^ SPCs is required for cold-induced BAT thermogenesis

BAT is the primary site of non-shivering thermogenesis, a major process of energy expenditure. Previous work has directly implicated Tregs and eosinophils in the induction of thermogenesis^19,25,30^, which prompted us to investigate whether the reduction in the frequency of BAT Tregs and eosinophils in SPC^IL33cKO^ mice would impact thermogenesis. Thermal imaging and temperature measurements of lean, unchallenged SPC^IL33cKO^ and IL33^fl/fl^ mice revealed no significant differences in BAT, core, or tail temperature (Figure 4A-C, Supplementary Figure 4A-B). Further corroborating this finding, no differences were detected in the energy expenditure and respiratory quotient of lean, unchallenged SPC^IL33cKO^ and IL33^fl/fl^ mice (Figure 4D, Supplementary Figure 4C-E), and qPCR analysis of BAT confirmed no significant differences in the expression of thermogenic genes between lean, unchallenged SPC^IL33cKO^ and IL33^fl/fl^ mice (Figure 4E). We next assessed BAT activity in cold-challenged, lean mice. We housed mice at thermoneutrality (30°C) for 10 days to reduce BAT thermogenesis, before subjecting one group to an additional 4°C cold challenge for 8h (Figure 4F). We first confirmed the robust, cold-induced upregulation of thermogenic genes such as *Ucp1, Pgc1a, Elovl3, Dio2, Lpl* and *Gpr3* between thermoneutral and cold challenged wildtype mice (Figure 4G), before assessing differences between cold-challenged IL33^fl/fl^ and SPC^IL33cKO^ mice (Figure 4H). Notably, SPC^IL33cKO^ mice were unable to upregulate both *Elovl3* and *Gpr3* following cold challenge (Figure 4I). *Elovl3* expression is highly correlated with recruitment of thermogenic brown adipocytes in BAT, and contributes to the production of <1% of the cellular BAT lipidome^31^. Additionally, *Gpr3* encodes a constitutively active, cold-induced G2-coupled receptor in brown and inducible beige adipocytes, upregulated in response to non-canonical lipolytic signals, and has been directly correlated with energy expenditure due to its constitutive activation^32^. Together, we show that loss of IL33 production by SPCs impairs adaptive BAT thermogenesis in metabolically challenged SPC^IL33cKO^ mice.

**Figure 4:**
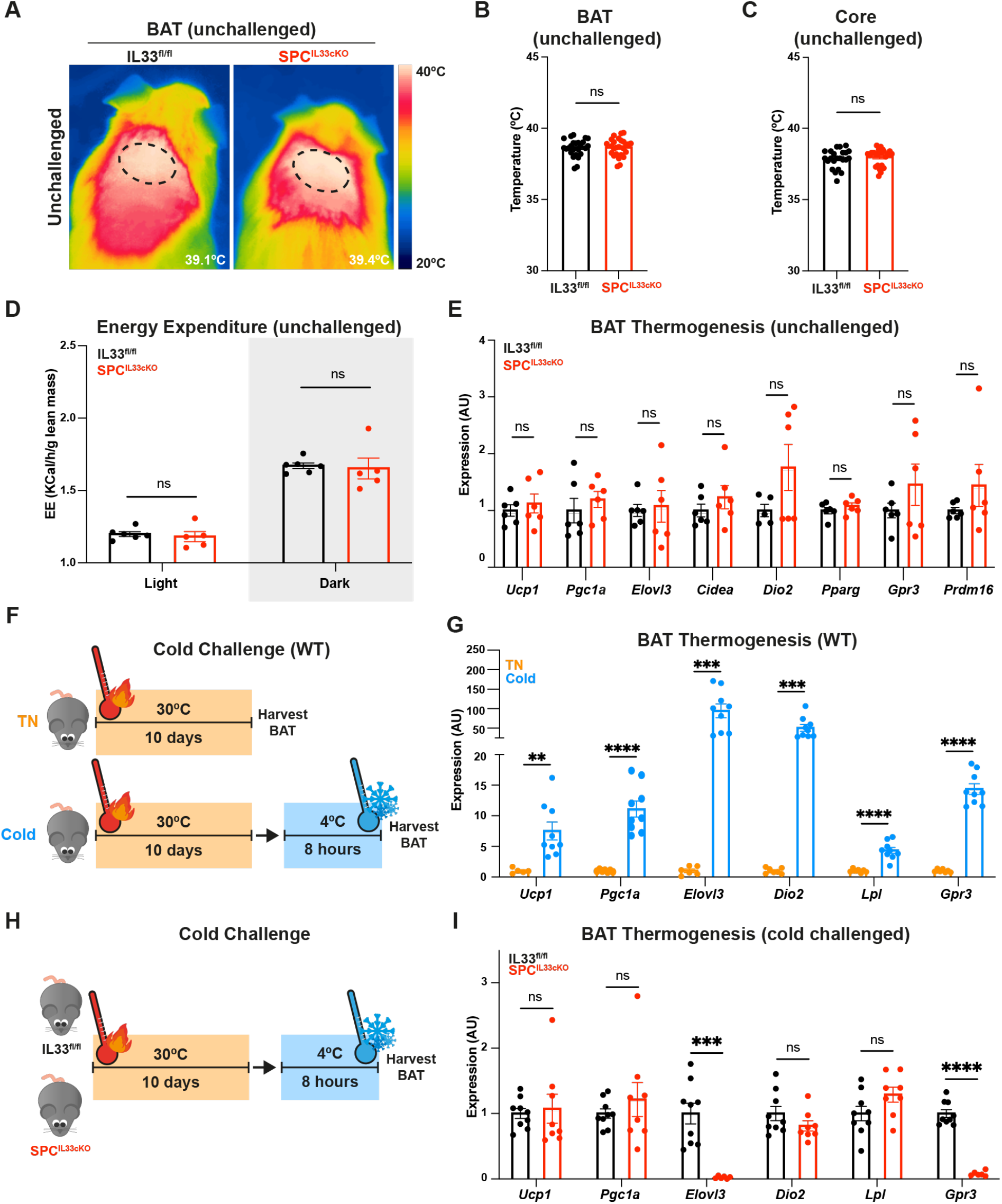
IL33 derived from LepR^+^ SPCs is required for cold-induced BAT thermogenesis. **A-B)** Brown adipose tissue (BAT) temperature is comparable in lean, unchallenged SPC^IL33cKO^ and IL33^fl/fl^ mice. Thermal imaging (**A**) and quantification (**B**) at room temperature (21°C). **A**) Dotted lines indicate region measured, bottom right = average temperature in image shown. **B)** Each data point represents average BAT temperature per mouse, calculated from <10 thermal images. n=25-27, 4 independent experiments. **C**) Core temperature of lean SPC^IL33cKO^ mice is unaltered at room temperature (21°C). n=22-24, 4 independent experiments. **D)** Energy Expenditure (EE) is comparable in lean SPC^IL33cKO^ and IL33^fl/fl^ mice. Each data point represents an average of EE per mouse during light/dark periods, measured over 5 days. EE normalised to lean mass. n=5-6, 1 independent experiment. **E)** qPCR of BAT from lean IL33^fl/fl^ and SPC^IL33cKO^ mice shows no difference in BAT thermogenic gene expression in unchallenged mice at room temperature (21°C). For each gene, data is presented relative to IL33^fl/fl^ expression level (=1). *Ucp1* = Uncoupled protein 1, *Pgc1a* = Peroxisome proliferator-activated receptor gamma coactivator 1-alpha, *Elovl3* = Elovl3fatty acid elongase 3, *Cidea* = Cell death inducing DFFA Like Effector A, *Dio2* = Iodothyronine Deiodinase 2, *Pparg* = Peroxisome proliferator activated receptor gamma, *Gpr3* = G-protein coupled receptor 3, *Prdm16* = PR/SET Domain 16. n=6, 2 independent experiments. **F)** Schematic representation: control WT mice were housed at thermoneutrality (30°C) for a period of 10 days before tissue harvest. Cold challenged WT mice underwent an additional 8-hour cold challenge (4°C). **G)** qPCR on the brown adipose tissue of wildtype thermoneutral (TN, orange) vs wildtype cold-challenged (Cold, blue) mice shows the robust upregulation of thermogenic gene expression following cold challenge. For each gene, data is presented relative to thermoneutral expression level (=1). *Ucp1* = Uncoupled protein 1, *Pgc1a* = Peroxisome proliferator-activated receptor gamma coactivator 1-alpha, *Elovl3* = Elovl3 fatty acid elongase 3, *Dio2* = Iodothyronine Deiodinase 2, *Lpl* = Lipoprotein Lipase, *Gpr3* = G-protein coupled receptor 3. n=6-9, 2 independent experiments. **H)** Schematic representation: cold-challenged IL33^fl/fl^ and SPC^IL33cKO^ mice were housed at thermoneutrality (30°C) for 10 days before an 8-hour cold challenge (4°C). **I)** qPCR of brown adipose tissue (BAT) shows impaired *Elovl3* and *Gpr3* upregulation in cold-challenged SPC^IL33cKO^ mice compared with cold-challenged IL33^fl/fl^ controls. For each gene data is presented relative to cold-challenged IL33^fl/fl^ expression level (=1). *Ucp1* = Uncoupled protein 1, *Pgc1a* = Peroxisome proliferator-activated receptor gamma coactivator 1-alpha, *Elovl3* = Elovl3 fatty acid elongase 3, *Dio2* = Iodothyronine Deiodinase 2, *Lpl* = Lipoprotein Lipase, *Gpr3* = G-protein coupled receptor 3. n=6, 2 independent experiments. Data presented as Mean+/-SEM. Student’s t-test, ns = non-significant, **p<0.01 ***p<0.001, ****p<0.0001.

### IL33 produced by SPCs is required for leptin’s rescue of metabolic adaptation to fasting

Finally, we developed a protocol to induce and subsequently rescue a state of metabolic adaptation to assess the physiological role of IL33 in SPCs. Metabolic adaptation is the reduction in energy expenditure in response to fasting to preserve energy stores, which we detect by the reduction of BAT temperature (Supplementary Figure 5A-B). In response to a 14h fast we found that expression of thermogenic genes *Ucp1, Elovl3* and *Dio2* was lower in SPC^IL33cKO^ BAT compared with IL33^fl/fl^ controls, indicating that SPC^IL33cKO^ mice exhibit a reduced capacity for thermogenesis with calorie restriction (Figure 5A). We used leptin to stimulate BAT thermogenesis, rescuing the fasting-induced metabolic adaptation (Figure 5B), and indeed observed that, relative to PBS-injected controls, leptin rescues the BAT temperature of fasted IL33^fl/fl^ mice (Figure 5C-D, Supplementary Figure 5C). However, in fasted SPC^IL33cKO^ mice, we observed that the BAT temperature is not rescued by IP leptin injection (Figure 5E-F, Supplementary Figure 5D), indicating that SPC^IL33cKO^ BAT is unresponsive to the leptin-induced stimulation of thermogenesis. Collectively, here we show that IL33 produced by SPCs is required for BAT response to a leptin challenge and metabolic adaptation to fasting.

**Figure 5:**
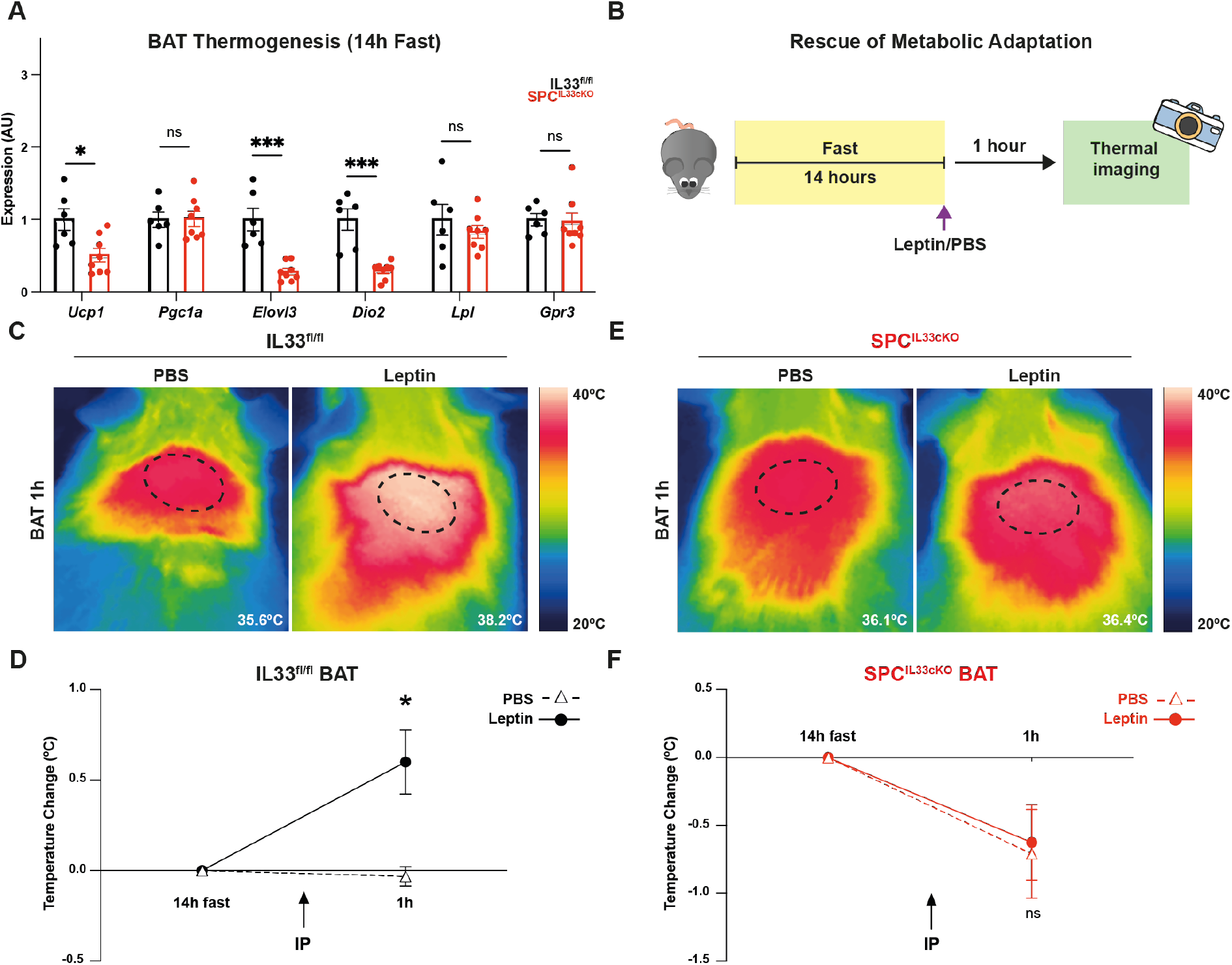
IL33 from SPCs mediates metabolic adaptation in BAT. **A)** Expression of *Elovl3* and *Dio2* thermogenic genes are reduced in the BAT of SPC^IL33cKO^ mice fasted for 14h, compared with 14h fasted IL33^fl/fl^ BAT. *Ucp1* = Uncoupled protein 1, *Pgc1a* = Peroxisome proliferator-activated receptor gamma coactivator 1-alpha, *Elovl3* = Elovl3 fatty acid elongase 3, *Dio2* = Iodothyronine Deiodinase 2, *Lpl* = Lipoprotein Lipase, *Gpr3* = G-protein coupled receptor 3. n=6-8, 2 independent experiments. **B)** Leptin-induced rescue of metabolic adaptation. Mice were fasted for 14 hours to induce metabolic adaptation before IP injection of Leptin or PBS (control). BAT temperature was measured 1 hour following IP injection. **C-F)** Leptin rescues metabolic adaptation in IL33^fl/fl^ mice (**C-D**) but not SPC^IL33cKO^ (**E-F**) mice. Representative thermal images of IL33^fl/fl^ BAT (**C**) and SPC^IL33cKO^ BAT (**E**) 1 hour following IP injection of either Leptin or PBS. Dotted lines indicate region from which the average BAT temperature was measured, bottom right = average temperature in thermal image shown. (**D, F**) BAT temperature was measured from mice injected with either Leptin (solid line) or PBS (dotted line). Presented relative to fasted BAT temperature. n=5-6, 2 independent experiments. Data presented as Mean+/-SEM. **A**) student’s t-test, **D, F**) 2-way ANOVA with Ŝidak correction for multiple comparisons. ns = non-significant, *p<0.05, ***p<0.001.

## Discussion

Obesity research has been divided into neuroendocrinology and immunometabolism, but a link is missing between these diverse fields of study. Here, we identify a LepR^+^, IL33-producing Sympathetic Perineurial Cell (SPC) barrier surrounding sympathetic ganglia and axon bundles in adipose tissues, providing a cellular link between these distinct fields. Across a range of different species, perineurial cells have been described as mesenchymal^33^, fibroblastic^34,35^, endothelial-like^36^ and epithelial^37^; with proposed functions including nerve protection, and regeneration^14–16,37^. We uncover unforeseen hormone-sensing and immunomodulatory roles for LepR+ SPCs, by showing that they are leptin-sensing and are capable of signalling BAT immune cells through the production of IL33.

The importance of IL33-producing stromal cells for immune recruitment and maintenance has recently been described in a variety of tissues. In fact, *Kuswanto et al*^38^ report an IL33^+^subset of perineurial cells in muscle, which based on our immunofluorescent imaging analysis are not LepR^+^ (Figure 2H), further indicating that LepR^+^ SPCs are a specialised subset of IL33-producing perineurial cells found in adipose tissues. Furthermore, two recent back-to-back papers have described the heterogeneity of IL33-producing mesenchymal cells in white adipose tissue, but did not identify an IL33-producing perineurial cell population^17,18^. We found that sympathetic axon bundles must first be dissected to isolate LepR^+^ SPCs, and thus propose that these barrier cells are a source of IL33 distinct from those previously reported by *Māhlakoiv et al*^17^ and *Spallanzani et al*^18^ using bulk analysis of adipose tissues. These studies involving either a total knockout of IL33^17^ or a PDGFRα-dependent conditional knockout of IL33 (PDGFRα^IL33cKO^)^18^ have been crucial in understanding the role of IL33 in adipose tissue, but have focussed solely on WAT and not BAT. A whole-body knockout of IL33 results in increased weight gain in the absence of metabolic challenge, whereas a PDGFRα-driven conditional deletion of IL33 was not sufficient to induce obesity in the unchallenged state, phenocopying SPC^IL33cKO^ mice (Figure 3C-G). Furthermore, we demonstrate that the frequency of visceral and subcutaneous WAT Tregs is unaltered in unchallenged SPC^IL33cKO^ mice, in comparison with the reduction observed in PDGFRα^IL33cKO^ visceral WAT^18^. Most notably, we show that deletion of *Il33* in SPCs results in a loss of BAT-resident anti-inflammatory Tregs and eosinophils, contributing to BAT inflammation (Figure 3I-K). IL33-mediated orchestration of immune cells by LepR^+^ SPCs is distinct in both mechanism and tissue from the GDNF-dependent immunosignalling reported in sympathetic associated stromal cells of the visceral WAT^39^, and further reinforces a specialised immunomodulatory role for perineurial cells.

We identify LepR^+^ SPCs as a significant contributor of IL33 to recruit/maintain a pool of BAT Treg and eosinophils, and corroborate previous reports suggesting a role in thermogenesis for these immune cells^19,25,30^. Although unchallenged SPC^IL33cKO^ mice have unaltered metabolism, dysfunction emerges upon metabolic challenge, with worsened HFD-induced obesity and impaired BAT thermogenesis in response to cold, fasting, and leptin stimulation. Collectively, we show that SPC-derived IL33 is critical for BAT response to metabolic challenge, establishing a role for LepR^+^ SPCs in the regulation of energy homeostasis. We uncover SPCs as an upstream, immunomodulatory, sympathetic stromal cell that is a missing link between leptin and immunometabolic mechanisms controlling body weight homeostasis, and obesity.

## Acknowledgements

We thank The University of Oxford Biomedical Services, The Don Mason Facility of Flow Cytometry (Dunn School, University of Oxford) and Light Microscopy Facility, Micron Advanced Bioimaging Unit (Dunn School, University of Oxford) for their help on this project. We would like to thank the Metabolic cage equipment facility Santiago and the Electron microscopy facility San Rafaelle, Ma’ayan Semo for help with animal welfare, and Cirenia Arias Baldrich (Cirenia Sketches) for illustrating diagrams for this project. We thank SIgN Immunogenomic core, A*STAR for helping to generate and analyse RNA sequencing data. This work has been supported by grants from Wellcome/HHMI International Research Scholar award 208576/Z/17/Z, ERC Consolidator grant – SympatimmunObesity ERC-2017 COG 771431, Wellcome Trust DPhil studentship awarded to E.H 222337/Z/21/Z, Ministerio de Ciencia y Universidades co-funded by the FEDER Program of EU (PID2021-128145NB-I00 and PDC2022-133958-I00) to M.L, Marie Curie Postdoctoral Fellowship FATARGET grant agreement no. 795891 awarded to N.M.S, British Heart Foundation DPhil studentship awarded to C.J.O.O, Novo Nordisk postdoctoral fellowship awarded to G.S and Pfizer’s 2022 Obesity Aspire Award.

## Author Contributions

A.I.D. conceptualised the study. B.A first visualised LepR^+^ SPCs, B.A, S.Y.Ö and N.M.S sorted LepR^+^ cells from SCG which were sequenced by S.C and F.G at SIgN Immunogenomic core, A*STAR. Transmission Electron Microscopy was performed by A.R and M.I on samples prepared by B.A. and N.M.S, S.Y.Ö created the first generations of SPC^IL33cKO^ mice. Flow cytometry was performed by E.R.H with the help of S.Y.Ö, C.J.O.O and C.Z. Sample preparation and confocal microscopy was performed by E.R.H, E.F and G.S. Analyses of published sc-seq data was performed by D.E.G and G.S. Metabolic phenotyping and subsequent analyses were performed by I.F.G and M.L. Weight acquisition (normal diet and high fat diet), fasting, cold challenge, leptin-induced rescue of metabolic adaptation and all accompanying thermal imaging and qPCR experiments were performed by E.R.H, G.S purchased and helped setup the cold chambers to meet H.O. standards. Blinded thermal imaging analyses were performed by E.R.H and E.F. E.R.H wrote the first draft of the manuscript and A.I.D the final version after several rounds of rewriting with E.R.H.

## Methods

### Animal husbandry

LepR^eYFP^ lineage-tracer and LepR^IL33KO^ mice were generated by crossing LepR^cre^ mice (Jax #008320) with either Rosa26^LSL-Chr2-eYFP^ (Jax #012569) or Il33^fl/fl^ (Jax # 030619) mice, respectively. Animals were bred and maintained at the University of Oxford under specific pathogen free conditions, on a 12h light/dark cycle at 21°C+/-1°C, 50% humidity +/-10%. Only age-matched male mice were used. Lean mice were 10-11 weeks old and unless otherwise stated (e.g., 14h fast) had *ad libitum* access to regular chow diet (normal diet, ND). All experiments were performed according to Institutional and UK Home Office regulations and the University of Santiago de Compostela Ethical Committee (15012/2020/010).

### Metabolic phenotyping

The metabolism of 10wo ND-fed IL33^fl/fl^ and SPC^IL33cKO^ was measured using indirect calorimetry (LabMaster, TSE Systems, Bad Homburg, Germany) as previously described^42^. This system is an open circuit instrument which determines O_2_ consumption (VO_2_), CO_2_ production (VCO_2_), Respiratory Quotient (RQ) - using the equation RQ=RER=VCO_2_/VO_2_ - and Energy Expenditure (EE). Mice were placed for 1 week prior to measurement for adaptation and measured over a period of 5 days. Nuclear Magnetic Resonance imaging (Whole Body Composition Analyser, EchoMRI) was used to measure body composition. Respiratory quotient and energy expenditure were normalised to lean mass.

### High fat diet challenge

At 8wo, mice were given *ad libitum* access to a high fat diet (60% fat, D12492: Research Diets) for a period of 12 weeks. Mouse body weight and food intake was measured throughout.

### Cold challenge

To reduce sympathetic activity onto BAT, 10wo IL33^fl/fl^ and SPC^IL33cKO^ mice were housed at thermoneutrality (30°C, HPP1060 Memmert climate chamber) for 10 days with *ad libitum* access to food and water. Thermoneutral control samples were collected, before experimental mice were housed at 4°C (HPP1060 Memmert climate chamber) for 8h and cold-challenged samples were collected. Thermoneutral and cold-challenged samples were collected at the same time of day, and immediately snap frozen. All mice were culled using CO_2_ (unperfused).

### Fasting and rescue of metabolic adaptation

To induce metabolic adaptation 10wo IL33^fl/fl^ and SPC^IL33cKO^ mice were fasted for 14h, with *ad libitum* access to water. Mice were then injected intraperitoneally (IP) with either 1 × PBS or 0.5mg/ml Leptin (10ul/g).

### Temperature measurements

BAT and tail temperature of unanaesthetised 10wo IL33^fl/fl^ and SPC^IL33cKO^ mice were measured using a thermal camera (Flir). Mice were shaved to expose the interscapular region >2 days prior to thermal imaging to avoid stress-induced BAT activity. Average temperature was calculated using Flir Tools software, where the average temperature was measured from a minimum of 10 images per mouse/ timepoint. All thermal image analysis was blinded. Core temperature was measured using a small rectal probe (Precision) on unanaesthetised mice.

### Immunofluorescence and confocal microscopy

10wo LepR^eYFP^ IL33^fl/fl^ and SPC^IL33cKO^ mice were given an IP overdose of pentobarbitone and perfused with 1x PBS, before dissection and fixation in 4% PFA (Thermofisher) overnight (4°C). Adipose tissue axon bundles were stained and imaged wholemount. For sectioned tissues (Figure 2, Supplementary Figure 2) samples were incubated in 30% sucrose (Sigma) overnight (for brains 5 days), before being embedded in OCT, snap frozen and cryosectioned (15 μm) using a Bright OTF5000 Cryostat. Wholemount and sectioned tissues were incubated in a blocking/permeabilisation buffer (3% BSA, 2% donkey serum, 1% Triton X-100, 0.1% Sodium Azide in 1x PBS) for 1h at room temperature. Overnight incubation with primary antibodies diluted in blocking/permeabilisation buffer (GFP – 1:1000 Ab13970 Abcam, TH – 1:1000 Ab152 Sigma, IL33 – 1:50 AF3626 R&D) were performed overnight at 4°C. Following washes with PBS, samples were incubated with secondary antibodies diluted 1:500 and DAPI (1:1000) in blocking/permeabilisation buffer (Dk-a-Ch-AF488 – 703-545-155 Jackson Immunoresearch, Dk-a-Rb-AF546 – A10040 Thermofisher, Dk-a-Gt-AF647 – A-21447 Life Tech) for 1h at room temperature. Samples were washed with PBS before being mounted with anti-fade medium (P36930, Invitrogen). Secondary only, DAPI-stained controls were used to ensure staining specificity. Z stack images were acquired using the LSM-880 confocal microscope.

### Transmission electron microscopy

10wo LepR^eYFP^ mice were given an IP overdose of pentobarbitone and perfused with 1x PBS. Axon bundles were dissected from the subcutaneous adipose tissue and fixed (4% PFA in 0.1M phosphate buffer, pH 7.4). Samples were washed with PBS before being incubated with glycine (50mM) and a permeabilisation solution (0.25% saponin, 0.1% BSA in 1x PBS). Adipose tissue axon bundles were then incubated with a blocking solution (0.2% BSA, 5% goat serum, 50 mM NH_4_Cl, 0.1% saponin, 150 mM NaCl in 20 mM phosphate buffer) for 20’, before incubation with anti-GFP antibody (1:500, A-11122 Invitrogen) for 2h at room temperature. Samples were washed with PBS and incubated with secondary antibody conjugated to 1.4nm gold particles (#2003 Nanogold; Nanoprobes). Nanogold particles were dilated with gold particle amplification solution (GE BEEM; Nanoprobes), according to the manufacturer’s instructions. Samples were wrapped with resin (Electron Microscopy Science, USA) and cured at 60°C for 48h. Resin blocks were sectioned using a Leica EM UC7 ultramicrotome, and ultra-thin slices (70-90 nm) were visualised using an FEI Talos 120 kV transmission electron microscope. Images were acquired using a Ceta 16M CMOS sensor camera (FEI, Netherlands).

### Single cell suspensions

For SPC phenotype, 10wo LepR^eYFP^ mice were given an IP overdose of pentobarbitone and perfused with 1x PBS before scWAT axon bundles, BAT axon bundles and Superior Cervical Ganglia were dissected. Axon bundles were digested for 30’ at 37°C (shaking) using 2.5mg/ml collagenase II (C6885, Sigma) and 4000U/ml hyaluronidase IV-S (H3884, Sigma) in Hank’s Balanced Salt Solution. Ganglia were digested using 2.5mg/ml collagenase I (C2674, Sigma) for 10’ at 37°C (shaking), before being washed with FACS buffer (2% FCS, 0.1% Sodium Azide in 1× PBS) and incubated in 0.25% trypsin for 30’ at 37°C (shaking). Axon bundles and ganglia were then mechanically digested using syringes with needles of decreasing size (23G, 25G, 27G) to create a single-cell suspension.

For immune cell phenotyping 10wo IL33^fl/fl^ and SPC^IL33cKO^ mice were culled using CO_2_ (unperfused), and spleens and adipose tissues were dissected. Spleens were homogenised through a 70μm filter. Adipose tissues were cut up, before digestion with 2.5mg/ml collagenase II (C6885, Sigma) for 30’ at 37°C (shaking).

### Flow cytometry

Single cell suspensions (prepared as above) were incubated with Fc block (553142, BD, 1:250) for 10’ at 4°C, followed by live/dead stain (Aqua – L34957, Invitrogen; Near IR – L34975, Invitrogen, Sytox Blue – S34857 Thermofisher, 1:250) for 30’ at 4°C. Samples were next stained with surface marker antibodies (CD45-AF700 1:500 103128 Biolegend, CD31-Pacific Blue 1:500 102421 Biolegend, TER119-BUV496 1:500 741079 BD, PDGFRα-APC 1:500 135907 Biolegend, ITGβ4-AF405 1:500 FAB405V Thermofisher, CDH5-PE-Cy7 1:500 138015 Biolegend, CD3e-PE-Cy5 1:250 100309 Biolegend, CD4-BV650 1:250 100545 Biolegend, CD8a-BV785 1:500 100749 Biolegend, CD25-ef450 1:200 48-0251-80 eBioscience, CD11b-ef450 1:500 48-0112-82 Thermofisher, CD64-PE 1:500 139303 Biolegend, Siglec F-BUV395 1:500 740280 BD) for 30’ at 4°C. Samples were then fixed using the FOXP3 fix/permeabilisation kit (00-5523-00, eBioscience) as per the manufacturer’s instructions; and subsequently incubated with antibodies for intracellular markers or corresponding isotype controls (IL33-PE 1:200 IC3626P R&D, FOXP3-PE 1:200 12-5773-82 Thermofisher, Isotype-PE 1:200 12-4321-83 Thermofisher) overnight at 4°C. Data was acquired using BD Fortessa X-20 cell analyser and analysed using FlowJo. Isotype controls or negative controls were used to set all gates.

### Bulk RNA-sequencing of LepR^+^ cells (SCG)

Single cell suspensions were prepared from the Superior Cervical Ganglia as above using 10wo LepR^eYFP^ mice. Live, LepR^eYFP+^ cells were sorted using a FACSAria IIu high-speed cell separator before cDNA libraries were prepared according to the Smart-Seq2 protocol ^43^ with the following adaptations: addition of 1mg/ml ultrapure BSA to lysis buffer, addition of 20μM template strand exchange nucleotide. cDNA library quality was checked CLS760672, CLS138948) and samples were subjected to paired indexing sequencing (Illumina HiSeq 4000). RNA sequencing data were analysed using Seurat.

### Analysis of published single-cell RNA sequencing data

*Lepr* and *Il33* co-expression in the hypothalamus and peripheral organs was assessed using data from ^20^ (GSE93374) and the *Tabula Muris* database ^21^ (GSE109774).

### RNA extraction, cDNA generation and qPCR

Frozen brown adipose tissue samples were homogenised in TRIzol (PreCellys 24 tissue homogeniser). After the addition of chloroform, the aqueous phase was carefully removed and nucleic acids were precipitated with isopropanol overnight at -20°C. Samples were DNAse treated (B0303s/ M0303s, NEB) and RNA concentrations were measured with the Nanodrop. cDNA was generated using 1000ng of RNA and SuperScript II Reverse Transcriptase (18064-014, Invitrogen) as per the manufacturer’s instructions. qPCR was performed with Power SYBR Green Master Mix (4367659, Thermofisher), 20ng of cDNA and primers (*Il1b*F-TGGACCTTCCAGGATGAGGACA, *Il1b*R-GTTCATCTCGGAGCCTGTAGTG, *iNos*F-GTTCTCAG CCCAACAATACAAGA, *iNos*R-GTGGACGGGTCGATGTCAC, *Tnfa*F-ATGAGCACAGAAAGCATGATC, *Tnfa*R-TACAGGCTTGTCACTCGAATT, *Il6*F-AAAGCCAGAGTCCTTCAGAGAGATAC, *Il6*R-CTGTTAG GAGAGCATTGGAAATTG, *Tbp*F-ACCGTGAATCTTGGCTGTAAAC, *Tbp*R-GCAGCAAATCGCTTGGG ATTA, *Ucp1*F-GTGAAGGTCAGAATGCAAGC, *Ucp1*R-AGGGCCCCCTTCATGAGGTC, *Pgc1a*F-CCCTGCCATTGTTAAGAC, *Pgc1a*R-TGCTGCTGTTCCTGTTTTC, *Elovl3*F-TTCTCACGCGGGTTAAAA ATGG, *Elovl3*R-GAGCAACAGATAGACGACCAC, *Cidea*F-TGCTCTTCTGTATCGCCCAGT, *Cidea*R-GCCGTGTTAAGGAATCTGCTG, *Dio2*F-CAGTGTGGTGCACGTCTCCAATC, *Dio2*R-TGAACCAAAGTT GACCACCAG, *Pparg*F-TCAAGGGTGCCAGTTTCG, *Pparg*R-GGAGGCCAGCATCGTGT, *Gpr3*F-ATCACCTGAGCAACCGAGAA, *Gpr3*R-AGATGGGGGTGCATTTTACA, *Prdm16*F-CAGCACGGTGAA GCCATT, *Prdm16*R-GCGTGCATCCGCTTGTG, *Lpl*F-CAGCTGGGCCTAACTTTGAG, *Lpl*R-CCTCTCTG CAATCACACGAA). Reactions were run using the BioRad CFX384 well plate reader (50°C: 2’; 95°C: 10’; **40x** 95°C: 15s, 60°C: 1’) with the addition of a final melt curve. All samples were loaded in technical triplicate. Singular melt curves were confirmed for each condition, and no-template controls were run for each gene to ensure no contamination. An average Ct value was calculated per sample, which was normalised against *Tbp* expression. Normalised expression levels were presented relative to the appropriate control (=1).

### Statistical analysis

Data are expressed as Mean +/-SEM. Statistical significance was determined by Student’s t test (2 groups) or ANOVA with post hoc two-tailed Bonferroni test when more than two groups were compared. p<0.05 was considered as significant.

