## Supplementary Figures for "Immunomodulatory Leptin Receptor^+^ Sympathetic Perineurial Cells Protect Against Obesity by Facilitating Neuroendocrine-Mediated Brown Adipose Tissue Thermogenesis"

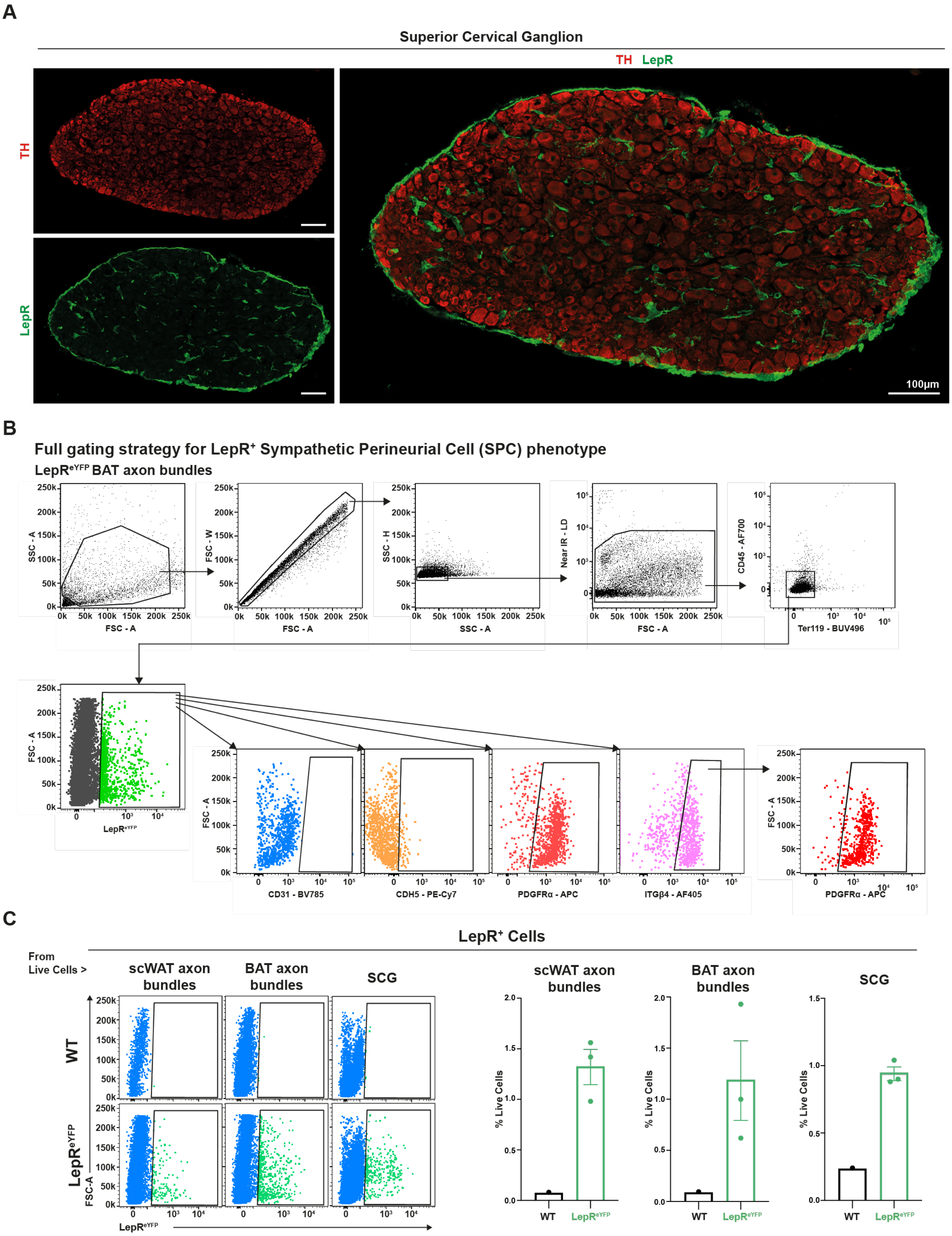


### Supplementary Figure 1

**A)** The Superior Cervical Ganglia (SCG) is surrounded by a LepR+ cell barrier (green). TH+ cell bodies (red). Scale bar = 100µm, cross section (20x). **B)** Full flow cytometric gating strategy for LepR+ Sympathetic-Associated Perineurial Cells (SPCs) phenotyping in sympathetic tissues. Most LepR+ cells in scWAT and BAT axon bundles are ITGβ4+PDGFRα+. **C)** LepR+ cells are present in sympathetic tissues. SCG and axon bundles dissected from the scWAT or BAT of LepReYFP mice (n=3).

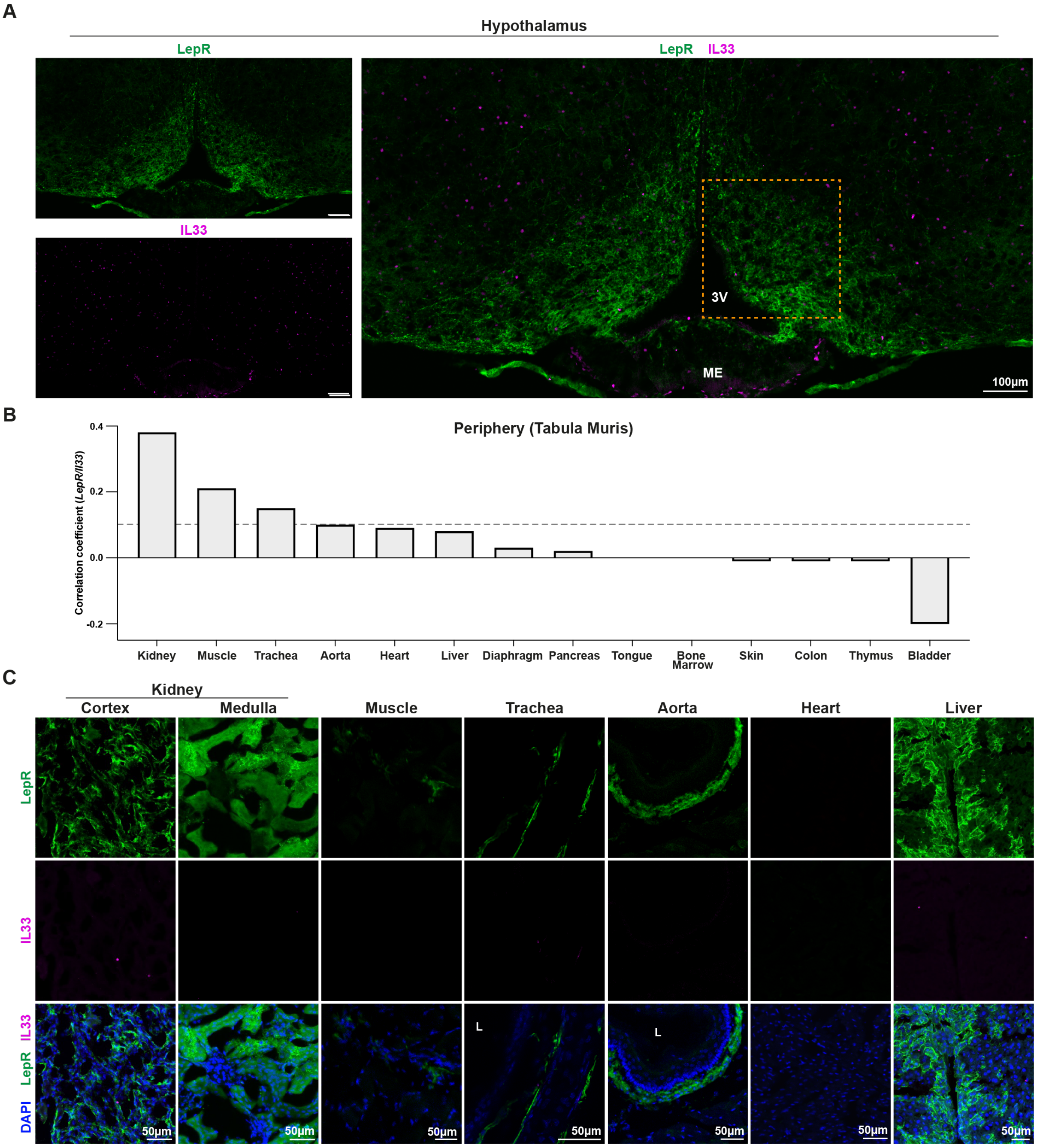


**Supplementary Figure 2**

**A)** Immunofluorescent staining of Coronal LepR$^{eYFP}$ hypothalamus section, maximum intensity projection image (10x). LepR$^{eYFP}$ (green), IL33 (magenta). Region marked with orange border is shown magnified in Figure 2G. 3V = third ventricle, ME = median eminence. Scale bar = 100µm. **B)** Co-expression of *Lepr* and *Il33* in peripheral mouse organs. Smart-Seq2 dataset, [21]. Grey dotted line marks correlation co-efficient of 0.1. Kidney, muscle, and trachea all have a correlation coefficient between 0.4-0.1. **C)** LepR$^{+}$IL33$^{+}$ cells were not detected in the kidney (medulla/ cortex), muscle, trachea, aorta, heart, and liver of lean LepR$^{eYFP}$ mice, 20x maximum intensity projection images. LepR$^{eYFP}$ (green), IL33 (magenta), DAPI (blue). Scale bars = 50µm. L = lumen.

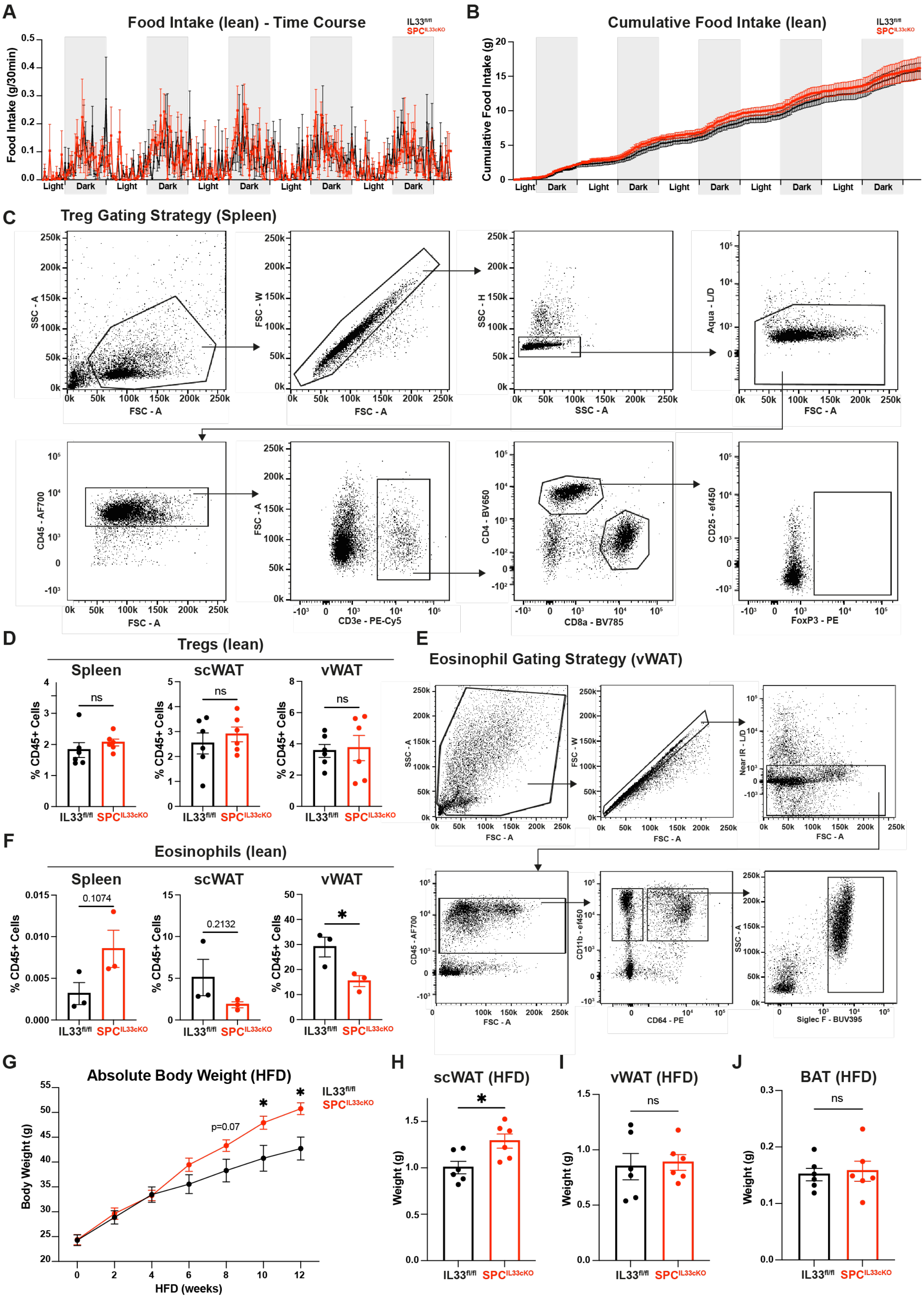


**Supplementary Figure 3**

**A-B)** Food intake (g/30min, **A**) and cumulative food intake (g, **B**) of lean (normal diet-fed) IL33$^{fl/fl}$ and SPC$^{IL33cKO}$ mice, measured over 5 days. n=5-6, 1 independent experiment. **C)** Flow cytometry regulatory T cell (Treg) gating strategy, stained with isotype control - PE. **D)** No differences observed in Treg frequency in the spleen, scWAT and vWAT of $SPC^{IL33cKO}$ mice. n=6, 2 independent experiments. **E)** Flow cytometry eosinophil gating strategy. **F)** No differences observed in eosinophil frequency in the spleen or scWAT of lean $SPC^{IL33cKO}$ mice. Eosinophil frequency is reduced in the vWAT of $SPC^{IL33cKO}$ mice. n=3, 1 independent experiment. **G)** Absolute body weight of $IL33^{fl/fl}$ and $SPC^{IL33cKO}$ mice fed high fat diet (HFD) from 8wo for 12 weeks. n=6, 2 independent experiments. **H-J)** $IL33^{fl/fl}$ and $SPC^{IL33cKO}$ mice fed HFD have comparable subcutaneous white adipose tissue (scWAT, **H**), visceral white adipose tissue (vWAT, **I**), and brown adipose tissue (BAT, **J**) weights. n=6, 2 independent experiments. Data presented as Mean+/-SEM. Student's t-test, ns = non-significant, *p<0.05.

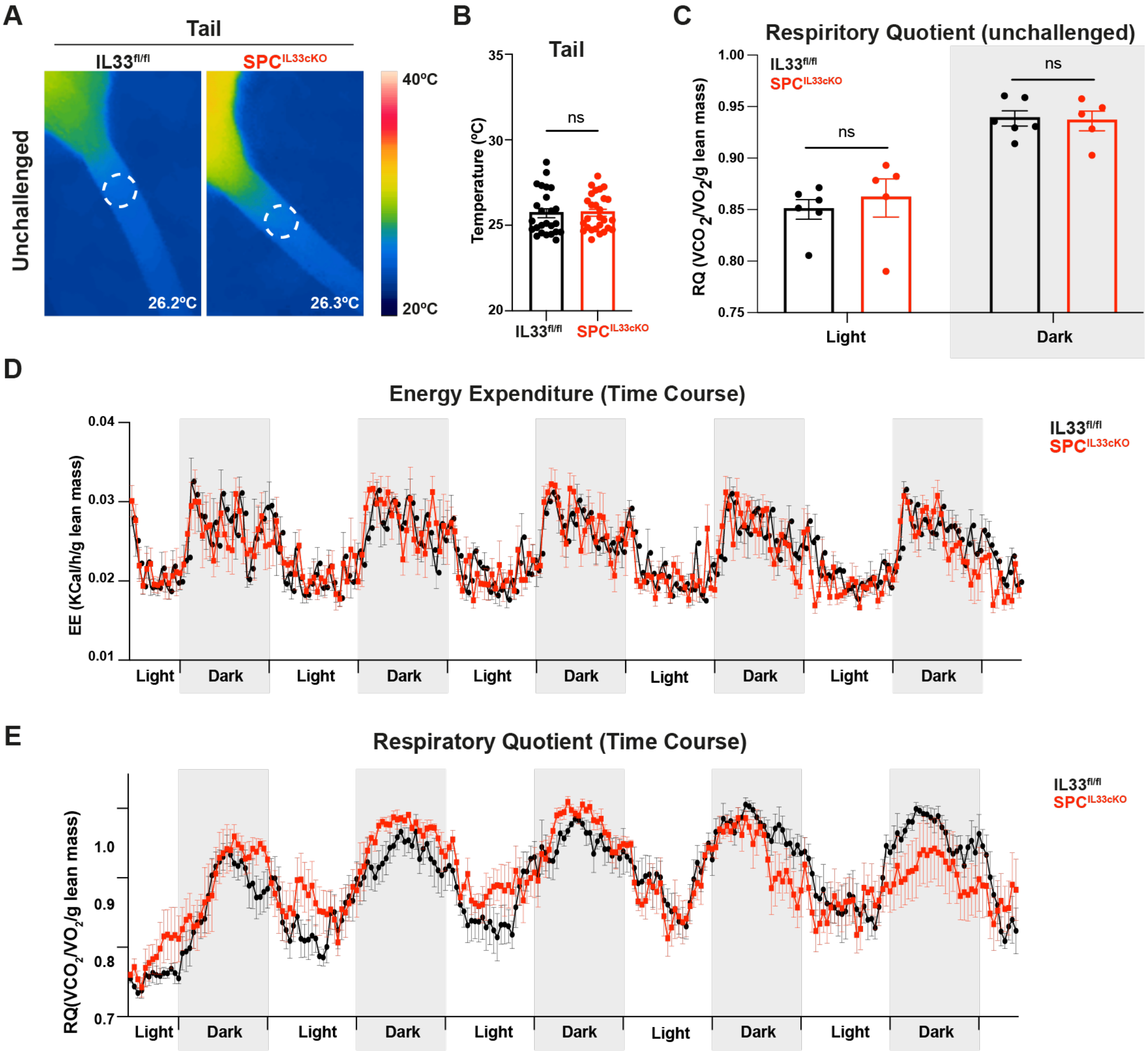


**Supplementary Figure 4**

**A-B)** IL33$^{fl/fl}$ and SPC$^{IL33cKO}$ mice have comparable tail temperatures. Representative thermal images (**A**) and quantification (**B**) of tail temperature at room temperature (21°C). **A**) Dotted lines indicate region from which the average temperature was measured, bottom right = average temperature in image shown. **B**) Each data point represents tail temperature from 1 mouse from >10 thermal images, n=25-27, 4 independent experiments. **C)** Respiratory Quotient (RQ) is unaltered in lean SPC$^{IL33cKO}$ mice. Each data point represents an average RQ per mouse during light/dark periods, measured over 5 days. RQ normalised to lean mass. n=5-6, 1 independent experiment. **D-E)** Energy Expenditure (EE, **D**) and Respiratory quotient (RQ, **E**) are comparable between IL33$^{fl/fl}$ and SPC$^{IL33cKO}$ mice, measured over 5 days. EE and RQ normalised to lean mass. n=5-6, 1 independent experiment. Data presented as Mean+/-SEM. Student's t-test, ns = non-significant.

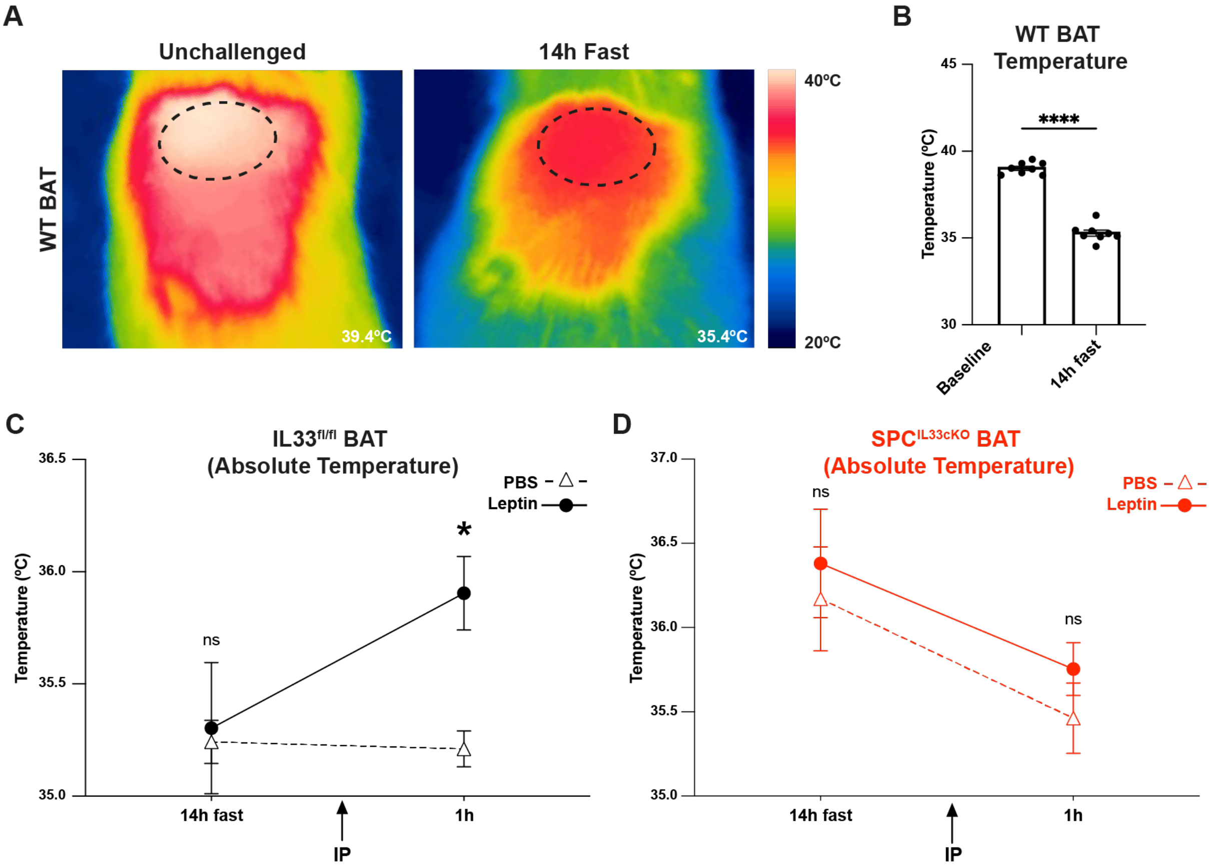


**Supplementary Figure 5**

**A-B)** Fasting induced metabolic adaptation. Lean, wildtype mice were fasted for 14 hours, when BAT temperature was measured. Thermal imaging (**A**) and quantification (**B**) at room temperature (21°C). **A**) Dotted lines indicate region measured, bottom right = average temperature in image shown. **B**) Each data point represents average BAT temperature per mouse, calculated from <10 thermal images. n=8, 2 independent experiments. **C-D)** Absolute IL33$^{fl/fl}$ (**C**) and SPC$^{IL33cKO}$ (**D**) BAT temperatures, measured from mice IP injected with either Leptin (solid line) or PBS (dotted line). n=5-6, 2 independent experiments. Data presented as Mean+/-SEM. B student's t-test, **C-D** 2-way ANOVA with Šidak correction for multiple comparisons. ns = non-significant, *p<0.05, **p<0.01, ***p<0.001, ****p<0.0001.
